# Evaluating the Efficacy of Serum-Free Media Supplemented with Protein Isolates for Bovine Satellite Cell Proliferation: A Sustainable Approach for Cultivated Meat Production

**DOI:** 10.1101/2024.08.23.609451

**Authors:** Arian Amirvaresi, Reza Ovissipour

**Author notes:** Corresponding author: Reza Ovissipour.

## Abstract

This study investigates the short- and long-term efficacy of serum-free media supplemented with protein isolates from algae, alfalfa, silkworm pupae, and grasshoppers for the proliferation of bovine satellite cells (BSCs). Fresh and spent media were analyzed to monitor metabolites, while cell proliferation was assessed using the CyQUANT assay. The results of the short-term growth study indicated that a lower concentration of alfalfa protein isolate (0.05 mg/mL) significantly enhanced cell proliferation, achieving a 1.47-fold increase compared to basal media, thereby demonstrating its potential as a viable alternative to fetal bovine serum (FBS). The capability of the developed serum-free media for cell passaging was examined using different coating strategies, with vitronectin outperforming others; however, it was unable to support the long-term growth of cells compared to FBS-containing media. Given the critical role of glutamine, this study also evaluates the impact of L-glutamine supplementation at different concentrations on cell growth and metabolism in alfalfa-based media. However, while glutamine supplementation showed trends toward increased cell growth, the enhancements were not statistically significant. Based on the results, this study highlights the potential of alfalfa protein isolate as a promising component of serum-free media for BSC proliferation and underscores the need for continued research into alternative protein sources and media formulations to support the sustainable and ethical production of cultivated meat.

## 1. Introduction

The cultivated meat industry is leading the path in food production innovation, offering a sustainable and advanced alternative that complements traditional animal agriculture. This burgeoning field aims to produce meat by cultivating animal cells *in vitro*, providing a more ethical and environmentally friendly option compared to conventional meat production (Dong et al., 2023; Srutee et al., 2022; Yang et al., 2023). One of the critical components in the production of cultivated meat is the culture medium used to grow animal cells such as bovine satellite cells (BSCs). Traditionally, Fetal Bovine Serum (FBS) has been the primary supplement in cell culture media due to its rich content of growth factors, hormones, and nutrients essential for cell growth and proliferation. FBS is known for its ability to support robust cell growth, but it presents significant ethical, environmental, and economic challenges. The supply of FBS is limited and subject to fluctuations in the livestock industry, making it an unsustainable option for large-scale cultivated meat production (Lezin et al., 2022; Reigado et al., 2024; Rossi, 2022). Consequently, there is an urgent need to develop serum-free media that can effectively replace FBS while supporting efficient and scalable cell growth. Serum-free media formulations have been a focus of intense research and development, and these formulations aim to provide all the necessary nutrients and growth factors required for cell culture without relying on animal-derived components (O’Neill et al., 2021).

Plant and insect protein isolates emerge as promising alternatives in this context (Batish et al., 2022; Timoneda et al., 2024; Reigado et al., 2024). These protein isolates offer a sustainable, ethical, and economically viable alternative to traditional serum supplements similar to FBS. Recent advancements in cell culture media have led to the development of formulations that include plant (Defendi-Cho & Gould, 2023), and insect protein hydrolysate (Batish et al., 2022). Several studies have focused on the development of serum-free media utilizing various protein isolations. Stout et al. (2022) introduced Beefy-9, a low-cost serum-free media derived from the B8 media originally designed for pluripotent stem cells. In their study, B8 supplemented with 0.8 mg/mL recombinant human albumin (Beefy-9), supports the short- and long-term expansion of BSCs while maintaining their proliferation and myogenicity (Stout et al., 2022). In another study, they presented Beefy-R, a novel serum-free medium where recombinant albumin (Beefy-9) was replaced with rapeseed protein isolate (RPI), derived from agricultural waste through alkali extraction. Beefy-R maintains BSC proliferation and myogenicity with improved growth rates compared to Beefy-9. Another study introduces a sustainable, serum-free, and grain-free culture medium for cultured meat production. Using nutrients from *Chlorella vulgaris* and mammalian cell growth factors, this media supports bovine myoblast proliferation without traditional fetal bovine serum and grain-derived nutrients. This approach reduces costs and enhances sustainability, providing an ethical alternative for cultured meat production (Yamanaka et al., 2023). Charlesworth et al. (2024) evaluated the use of hydrolysates from Kikuyu grass, Alfalfa grass, and cattle rearing pellets as partial replacements for FBS in the cultivation of C2C12 myoblast cells for cultivated meat production. The findings showed that while these hydrolysates alone were less effective under serum-free conditions, their growth-promoting effects were significantly improved when combined with minimal serum (0.1%) and further enhanced with insulin, transferrin, and selenium (ITS). Cattle rearing pellet hydrolysates exhibited the most substantial growth promotion, followed by Kikuyu and Alfalfa (Charlesworth et al., 2024).

In our previous studies, hydrolysates from black soldier fly larvae (BSFL), crickets, oysters, mussels, and lugworms have been evaluated for their effectiveness in supporting cell growth. These protein sources provide high-quality, essential amino acids and exhibit suitable functional properties (Batish et al., 2022). Recently, we developed a method for bioconverting BSFL into cultivated meat media components using the blue catfish gut microbiome. Through this study, not only different protein sources were used, but also novel processing methods such as gut biome assisted fermentation were applied (Timoneda et al., 2024). Another study by our research group investigates plant- and microbial-derived protein hydrolysates as alternatives to FBS in zebrafish embryonic stem cell culture media. Our findings reveal that while high concentrations of hydrolysates (1-10 mg/mL) inhibit cell growth, lower concentrations (0.001-0.1 mg/mL) enhance cell proliferation and viability, particularly when combined with reduced serum levels (1-2.5%) (Amirvaresi & Ovissipour, 2024). Therefore, exploring other plant-derived proteins or novel protein sources holds significant potential, offering new insights for the development of innovative serum-free media.

This research aims to explore the effects of protein isolates from algae, alfalfa, silkworm pupae, and grasshopper on the proliferation of bovine satellite cells (BSCs) in serum-free media, presenting a cost-effective and environmentally sustainable alternative. The primary objectives of this study are to compare the impacts of these protein isolates on BSC proliferation, determine the optimal concentrations for each isolate, analyze their amino acid and protein compositions, and evaluate metabolic changes in both fresh and spent media.

## 2. Materials and methods

### 2.1 Culture and maintenance of bovine satellite cells (BSC)

Bovine Satellite Cells (BSCs) were obtained from Professor Kaplan’s lab at Tufts University and were cultured under optimized conditions to facilitate their proliferation and maintenance. The cells were cultured in an ordinary growth medium (OGM) consisting of Dulbecco’s Modified Eagle Medium (DMEM) (Thermo Fisher, #10566024), supplemented with 10% fetal bovine serum (Thermo Fisher, #26140079), 1 ng/mL human Fibroblast Growth Factor-2 (FGF-2) (Thermo Fisher, #100-18B), and 1% antibiotic-antimycotic solution (Thermo Fisher, #15240062). Culturing flasks were coated with 0.1% (W/V) gelatin (VWR, #97062-618). For passaging, the cells were grown to around 70-80% confluency and detached with a 0.25% trypsin-EDTA solution (Thermo Fisher, #25200056). At all stages, the cells were maintained at 37°C and 5% CO_2_.

### 2.2 Protein extraction

Protein isolates were extracted through an acid-base precipitation method (Hojilla-Evangelista et al., 2017) with slight modifications. Initially, 100 grams of each source material—algae (chlorella, Micro Ingredients), alfalfa (Viking Farmer), silkworm pupae (Chubby), and grasshopper (Fluker’s)—was ground to a fine powder. This was followed by alkaline treatment with 1L of 0.1 M NaOH, adjusting the pH to 11 for protein solubilization. The mixture was sonicated for 10 minutes and stirred for 2 hours at 50°C. After extraction, the mixture was centrifuged at 4000*g* (AccuSpin MAX+ R 4 L Refrigerated Centrifuge) for 15 min to separate the supernatant containing solubilized proteins. The pH of the supernatant was then adjusted to 3.5-4.5 (isoelectric point, depends on protein) using HCl to precipitate the proteins. Following a second centrifugation step (4000*g*, 20 min), the protein-enriched pellet was collected, freeze-dried, and kept in a −20°C freezer. Quantification of the isolated proteins was achieved through the Bradford method based on the manufacturer protocol (Figure S1).

### 2.3 Protein isolate short-term growth study

The growth of bovine muscle cells in a serum-free media was evaluated over a period of three days. Briefly, a 96-well plate (VWR, #734-2327) was coated with a 1% w/v gelatin solution and incubated for 30 min at 37°C. The cells were seeded at a density of 8000 cells per well in serum-containing medium and incubated to adhere overnight. The following day, the serum-containing medium was removed, and the cells were rinsed with DPBS. After aspirating the DPBS, fresh media containing HiDef (Defined Bioscience, #LSM-102-6), along with HiDef supplemented with protein isolates (PIs) at concentrations of 0.05, 0.25, and 0.5 mg/mL, was added to cells. The cells were then incubated at 37°C with 5% CO_2_ for three days. Each experimental condition was replicated three times per plate, with two plates prepared for further analysis. On the third day, the cells were imaged using an inverted microscope (CKX53 Olympus).

Furthermore, cell proliferation was assessed using the CyQUANT™ NF cell proliferation assay kit (Thermo Fisher, #C35006), following the manufacturer’s protocol. To ensure accuracy and reliability, the assay was conducted in triplicate.

### 2.4 Live/dead cell viability assay

Cell viability was assessed using the LIVE/DEAD Cell Imaging Kit (Thermo Fisher, #R37601) and based on the manufacturer’s protocol. BSCs in OGM, Hidef, and alfalfa were incubated with calcein AM, which stains live cells green, and BOBO-3 Iodide, which stains dead cells red, for 30 minutes at room temperature. After incubation, cells were washed with PBS and imaged using the fluorescence microscope.

### 2.5 Immunostaining analysis

On the third day, bovine satellite cells cultured in ordinary growth media, HiDef B8 media, and alfalfa protein-supplemented media were assessed for cell proliferation and differentiation potential using a PAX7 immunostaining assay. Cells were fixed with 4% paraformaldehyde (Thermo Fisher, #J61899.AP) for 30 minutes at room temperature, followed by permeabilization with 0.5% Triton X-100 (Millipore Sigma, #T8787) for 20 minutes. The cells were then blocked with 5% bovine serum albumin (BSA) (Thermo Fisher, #16210064) for 1 hour to prevent nonspecific binding. Primary antibody incubation was performed using anti-PAX7 (1:100 dilution, Thermo Fisher, #PA5-68506) overnight at 4°C. After washing with PBS, cells were incubated with secondary anti-body (anti-rabbit (1:500 in blocking buffer, Thermo Fisher, #A-11072, 1:500 in blocking buffer)) for 1 hour at room temperature in the dark. Nuclei were counterstained with DAPI (1:100 diltuin, Abcam, #ab104139) for 15 minutes. The cells were then washed, and fluorescence images were captured using the microscope.

### 2.6 Protein isolate passaging experiment

The protein derived from alfalfa demonstrating superior performance in the short-term growth study was selected for subsequent passage experiments. In experiment I, 6-well plates were prepared in triplicate by coating with a 0.1% gelatin solution, followed by incubation at 37°C with 5% CO_2_ for 30 minutes. Subsequently, 25,000 cells per well were seeded in OGM and incubated overnight to facilitate cell attachment. Following incubation, wells were washed with PBS, and then OGM, HiDef B8 media, and alfalfa protein-supplemented media were added, with each condition tested in triplicate. Upon reaching approximately 70% confluency, cells were detached using 0.25% trypsin-EDTA solution for the OGM condition (5 minutes) and TrypLE Express (Thermo Fisher, #12604021) for the serum-free media condition (10-15 minutes). The cells were then diluted with the appropriate serum-free or serum-containing media, centrifuged at 400 *g* for 5 minutes, counted (Thermo Fisher, Countess™ 3 FL automated cell counter), and seeded at a density of 25,000 cells per well for the next passage. Cell growth and proliferation were monitored over the passage steps using cell counting and microscopic images.

To assess the effect of different coating materials on cell attachment, growth, and proliferation, a parallel experiment (experiment II) was conducted using 6-well plates. The wells were coated with 1.5 µg/cm² (Stout et al., 2023) vitronectin (Thermo Fisher, #A14700), fibronectin (R&D Systems, #ACFP4305B), or laminin (Iwai North America, #N-892012), with each coating condition tested in triplicate. The protocol for seeding and media changes was similar to that of the gelatin coating experiment. This experimental design enabled the evaluation of the influence of various coating conditions on cell behavior across multiple passages.

### 2.7 Fresh and spent media analysis

Fresh and spent culture media were analyzed using the BioProfile Flex2 Analyzer (Nova Biomedical, USA) to assess glucose, lactate, glutamine, glutamate, and ammonium levels, thereby evaluating metabolic activity and nutrient consumption.

### 2.8 Glutamine experiment

A short-term experiment (based on section 2.3) was conducted using HiDef basal media and alfalfa protein isolate media to assess the impact of glutamine on cell proliferation. These media were supplemented with L-glutamine at concentrations of 0.5, 1.6, and 3.0 mM. Cells were cultured in triplicate for each condition, including controls without glutamine supplementation, and incubated for three days. On the third day, fresh and spent media were analyzed for levels of glutamine, glutamic acid, glucose, lactate, and ammonium. Cell proliferation was evaluated using the CyQUANT assay, with growth measured relative to the basal media.

### 2.9 Amino acid analysis

A known quantity of standard solution was introduced to a tube containing internal standards, dried using a vacuum concentrator (SpeedVac), and accurately weighed. Each tube was filled with 300 µL of 6N hydrochloric acid, sealed under an argon atmosphere, and hydrolyzed at 110°C for 22 hours. Post-hydrolysis, the samples were centrifuged to remove debris, and a 40 µL aliquot of the supernatant was dried under vacuum. The dried samples were reconstituted with 0.4N borate buffer and analyzed using an Agilent Infinity 1260 HPLC system with a specified mobile phase gradient for maximum resolution and sensitivity.

### 2.10 Statistical analysis

Statistical analyses and graphical representations were conducted using GraphPad Prism 10.0 software (San Diego, CA, USA). Comparative significance between two groups was evaluated via a t-test, with statistical significance annotated for each dataset.

## 3 Results and discussion

### 3.1 Protein extracts and amino acids compositions

Proteins were isolated from alfalfa, algae, grasshopper, and silkworm pupae using an acid-base extraction method with an isoelectric point range of 3.5-4.5. The extraction yields for alfalfa, algae, grasshopper, and silkworm pupae were 2.8%, 9.7%, 28.8%, and 13.5%, respectively. However, further optimization of the extraction protocol can significantly enhance protein yields. The economic and environmental advantages of these protein isolates make them excellent candidates for developing serum-free media. Specifically, alfalfa protein is a particularly promising candidate due to its lower production costs compared to traditional animal-based proteins. Alfalfa is a highly sustainable crop, characterized by its ability to fix atmospheric nitrogen, thereby reducing the need for synthetic fertilizers and use water source efficiency, making it an environmentally friendly choice. These sustainable agricultural practices, combined with the high nutritional value of alfalfa protein, ensure a reliable, cost-effective, and eco-friendly supply of high-quality protein for serum-free media. This supports the advancement of sustainable and ethical cell culture practices (Latif et al., 2023). Amino acid analysis of proteins was conducted to identify trends and concentrations of essential and non-essential amino acids in protein isolate, finding potential patterns or specific amino acids that could influence cell growth. Based on the amino acid results (Table 1) and the cell proliferation findings, the quality and composition of amino acids present in alfalfa were more aligned with the nutritional needs of bovine satellite cells for growth and proliferation. This indicates that these cells require specific amino acids at precise concentrations for optimal performance, rather than simply increasing amino acid quantity.

**Table 1:** Amino acid analysis for four protein isolates including pupae, grasshopper, alfalfa, and algae.

| *mg of AA/g of protein* |  |  |  |  |
| --- | --- | --- | --- | --- |
| *Amino acid* | *Pupae* | *Grasshopper* | *Alfalfa* | *Algae* |
| *ASX* | 12.24 | 11.00 | 9.91 | 9.75 |
| *GLX* | 13.01 | 15.94 | 13.00 | 13.53 |
| *SER* | 5.80 | 6.40 | 5.02 | 5.04 |
| *HIS* | 2.66 | 2.60 | 2.24 | 1.88 |
| *GLY* | 4.53 | 4.69 | 5.19 | 5.86 |
| *THR* | 4.51 | 4.89 | 5.22 | 5.19 |
| *ALA* | 2.49 | 2.92 | 1.85 | 1.57 |
| *ARG* | 12.45 | 13.80 | 15.30 | 20.90 |
| *TYR* | 8.52 | 6.45 | 4.26 | 4.10 |
| *VAL* | 6.09 | 5.87 | 6.43 | 6.35 |
| *MET* | 3.22 | 2.38 | 2.00 | 2.50 |
| *PHE* | 6.80 | 5.07 | 6.01 | 5.65 |
| *ILE* | 5.34 | 5.71 | 5.12 | 4.41 |
| *LEU* | 8.01 | 8.69 | 7.97 | 8.70 |
| *LYS* | 3.18 | 3.08 | 4.57 | 3.07 |
| *PRO* | 1.21 | 0.50 | 5.90 | 1.51 |

ASX: Aspartic Acid or Asparagine, GLX: Glutamin and Glutamic acid, SER: Serine, HIS: Histidine, GLY: Glycine, THR: Threonine, ALA: Alanine, ARG: Arginine, TYR: Tyrosine, VAL: Valine, MET: Methionine, PHE: Phenylalanine, ILE: Isoleucine, LEU: Leucine, LYS: Lysine, PRO: proline, AA: Amino acid

Alfalfa’s amino acid composition, especially the high concentrations of Arginine (ARG) and notable amounts of Proline (PRO), could be particularly beneficial. Arginine is essential for protein synthesis and cell replication, while Proline supports collagen synthesis and extracellular matrix formation, crucial for satellite cell function (O’Neill et al., 2021). Furthermore, the improvement observed with the lowest alfalfa concentration might also reflect an optimal nutrient density that avoids the potential negative effects of high concentrations, such as osmotic stress or metabolic overload, which could inhibit cell growth or function. It is worth noting that it is important to establish a baseline of the amino acid content. This baseline will provide insight into the initial availability of essential and non-essential amino acids and help determine whether the initial concentrations align with the specific needs of the cells.

### 3.2 Short-term assessment of protein isolates: effects on cell growth and morphology

In the short-term, three-day experiment, the effects of four distinct protein isolates—algae, alfalfa, silkworm pupae, and grasshopper—at concentrations of 0.5, 0.25, and 0.05 mg/mL on cell growth and morphology of bovine satellite cells were investigated. On the third day, microscopic analysis revealed significant variations in cell growth and morphological characteristics across the different protein treatments (Figure S2). None of the tested concentrations of algae, grasshopper, or silkworm pupae proteins supported cell growth; instead, they induced apoptosis at all levels. However, for alfalfa, only higher concentrations (0.5 mg/mL and 0.25 mg/mL) of the protein showed cytotoxicity. Additionally, the cellular morphology under these conditions was suboptimal (Figure S2). Cell growth inhibition, especially at higher concentrations of protein, caused rapid cell death, suggesting the presence of additional growth-inhibiting or cytotoxic components within them. These components likely hinder cell growth when their concentration surpasses a certain threshold. Potential reasons for this could include impurities from the extraction process or naturally occurring substances that become toxic at higher concentrations, protein aggregation disrupting cellular processes, metabolic overload leading to the accumulation of toxic metabolites, and unknown bioactive compounds that interfere with cell signaling pathways or induce apoptosis (Hogwood et al., 2013; Stout et al., 2023). These results were in agreement with other researchers’ findings. For example, Batish et al. (2022) used Black soldier fly, crickets, oysters, mussels, and lugworms at 0.001–10 mg/mL concentrations and reported cytotoxicity of these proteins at high concentrations (Batish et al., 2022). Cytotoxicity of high concentrations of pea, mushroom, yeast, and algae protein hydrolysates has also been reported on zebrafish cell lines (Amirvaresi & Ovissipour, 2024). These factors highlight the importance of optimizing protein concentrations in cell culture media to avoid inhibitory effects and ensure optimal cell growth. In contrast, alfalfa protein at the lowest concentration (0.05 mg/mL) exhibited significant positive effects on both cell growth and morphology, closely aligning with the outcomes observed with ordinary growth medium (20% FBS) (Figure 1a-c) and Figure S2). Recently, the positive impacts of alfalfa protein hydrolysates on C2C12 cell growth were reported (Charlesworth et al., 2024). It was shown that alfalfa hydrolysates can promote cell growth only in the presence of low concentrations of serum (0.1%) or insulin, transferrin, and selenium (Charlesworth et al., 2024).

**Figure 1.**
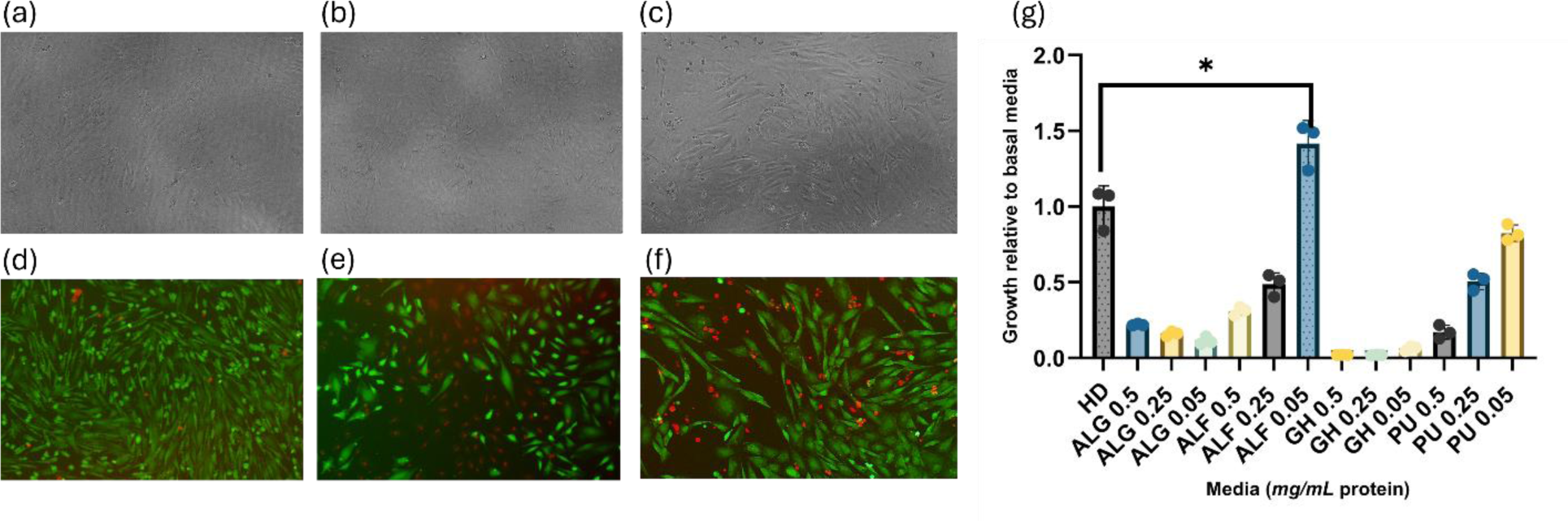
Evaluation of Cell Viability and Growth in Various Culture Conditions. (a-f) Brightfield and Live/Dead cell viability assays under different media conditions. Brightfield images: (a) OGM (20% FBS), (b) 0.05 mg/mL alfalfa protein, and (c) HiDef media. Alfalfa protein at the lowest concentration (0.05 mg/mL) demonstrated significant positive effects on cell growth and morphology, comparable to HiDef media as the control group. (d), (e), and (f) illustrate the results of the live/dead cell viability assay: (d) OGM (20% FBS) (high viability, minimal cell death, healthy morphology), (e) HiDef B8 media (moderate viability, notable cell death, cells show shrinkage), and (f) alfalfa-supplemented media (high viability, minimal cell death, healthy morphology similar to 20% FBS media). (g) CyQUANT assay results from a three-day cell proliferation study conducted to evaluate cell growth in media supplemented with different protein isolates compared to HiDef (HD) media. The assay results show cell growth in media containing protein isolates from algae (ALG), alfalfa (ALF), silkworm pupae (PU), and grasshopper (GH) at concentrations of 0.5, 0.25, and 0.05 mg/mL. The data specifically highlight that only alfalfa protein at the lowest concentration (0.05 mg/mL) significantly improved cell proliferation and population compared to the HiDef control. Statistical significance was determined using an unpaired t-test between the control (HiDef) and alfalfa 0.05 mg/mL, with a significance of p < 0.0251 (*). Results are based on n=3 distinct samples. The scale of each image is 100 μm.

The live/dead staining, as demonstrated in Figure 1(d-f), provides insights into the cell viability and cellular response under varying culture conditions. In Figure 1d, cells cultured in ordinary growth media (20% FBS) exhibit predominantly green fluorescence, indicating high cell viability with minimal red fluorescence, suggestive of a low incidence of cell death. Moreover, cell morphology appears normal, and the cells exhibit an elongated shape. Conversely, Figure 1e shows cells maintained in Hidef B8 media, where a more balanced distribution of green and red fluorescence is observed. This indicates a moderate level of cell mortality and a significant number of compromised cells. Moreover, the cell morphology in Hidef B8 media is notably altered, with evident spindle-shaped cells and signs of shrinkage that might indicate the onset of cell death, compared to those in FBS-supplemented media. Cells cultured in alfalfa-supplemented media (Figure 1f) demonstrate normal morphology, closely resembling that of the ordinary growth media condition. The prominent green fluorescence in this group suggests a high viability, similar to the cells in OGM containing 20% FBS.

CyQUANT assay was conducted to evaluate cell proliferation. An unpaired *t-test* determined the significance between HiDef and alfalfa media. Alfalfa significantly enhanced cell proliferation and population, achieving a 1.47-fold increase compared to HiDef, thereby demonstrating its superior efficacy in promoting cell growth (Figure 1g). The findings from the CyQUANT assay, combined with the microscopic and viability analyses, point to alfalfa protein as a promising, cost-effective, and sustainable option for enhancing cell culture media. Alfalfa effectiveness at low concentrations, along with its ability to maintain cell viability and morphology comparable to standard growth media, positions alfalfa protein as a potentially superior supplement for promoting cell growth in various research and biotechnological applications.

Immunostaining was performed (on day three) to evaluate the phenotype of proliferative bovine satellite cells (BSCs) in OGM, HiDef, and alfalfa media. DAPI, a cell nucleus staining marker, showed consistent results across all media types, displaying comparable basic cellular health and proliferation rates. Additionally, PAX7 staining was positive in all media types, which confirms the maintenance of satellite cell identity and myogenic potential. These findings suggest that the serum-free media used in this study did not significantly affect the cells proliferative capacity and myogenic potential, indicating that all the media tested are adequate for maintaining these key characteristics in culture.

**Figure 2.**
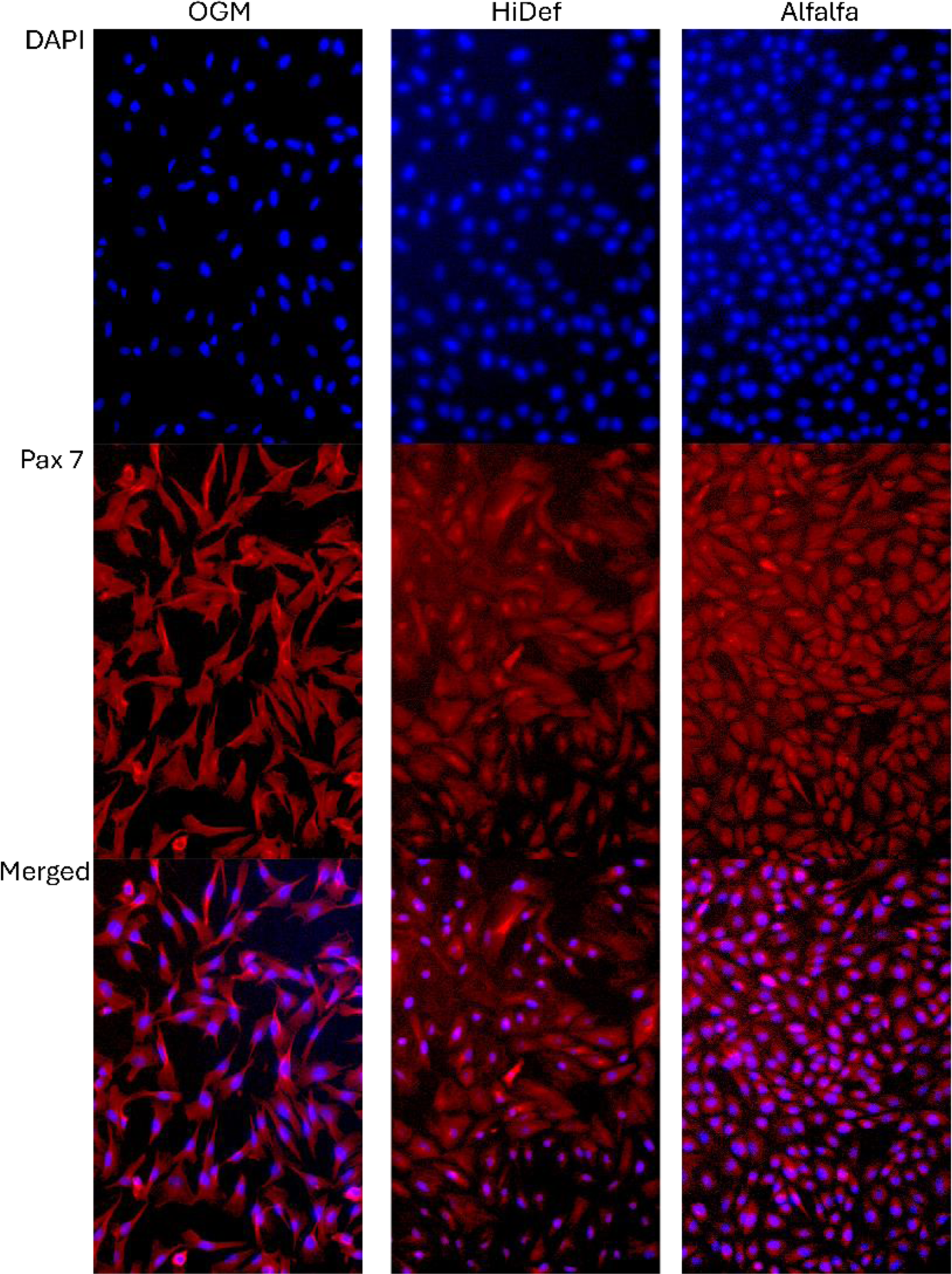
DAPI and Pax 7 immunostaining of proliferative cells cultured in OGM, HiDef, and alfalfa-supplemented media. Consistent results were observed across all media types. The scale of each image is 100 μm.

Based on these results, one possible reason for the positive effects of alfalfa protein on cell growth, morphology, and proliferation in serum-free media could be largely attributed to its higher albumin content, which is notably present in much lower quantities in other plant proteins (Altomare et al., 2020; Guarnieri et al., 2013; Hadidi et al., 2023). Albumin plays a crucial role in mammalian cell culture by transporting key molecules such as fatty acids and hormones into cells, supporting cell growth and survival, and offering protection against oxidative stress and mechanical damage in bioreactors (Francis, 2010). This aligns with the other researcher’s findings. For example, Stout et al. (2022), observed that the addition of recombinant albumin to cell culture media significantly enhanced cell growth, maintained cellular morphology, and promoted proliferation (Stout et al., 2022). Additionally, plant proteins like rapeseed, which have a high albumin content, could potentially replace recombinant albumin in serum-free media for cultivated meat (Stout et al., 2023). Similarly, Messmer et al. (2022) underscored the importance of albumin in serum-free media formulations, noting that while albumin alone did not markedly increase the fusion index of cells into multinucleated myotubes, its combination with other factors like insulin and transferrin improved differentiation outcomes (Messmer et al., 2022). Plant-based materials with high albumin content, previously used for cell proliferation (Yamane et al., 1981a,b; Georg et al., 2009), were compared to rapeseed protein isolates (Stout et al., 2023). None of these materials indicated effectiveness at the tested concentrations, suggesting that albumin concentration alone may not be the only key factor for cell growth. It is likely that a combination of multiple factors influences cell growth promotion.

### 3.3 Enhanced cell adhesion and proliferation through different coating strategies

A multi-passage experiment was conducted to evaluate the growth characteristics of BSCs in various media formulations, including OGM, HiDef, and alfalfa-supplemented media. Initially, 1 mL of 0.1% gelatin was used to coat all wells of the 6-well plates in experiment I. By day three, when cell confluency reached around 60-70%, cells were counted, and the doubling times were calculated for each media. The calculated cell doubling times were 14 hours for OGM, 32 hours for HiDef, and 27 hours for the alfalfa-supplemented media. Subsequently, cells were prepared for the next seeding phase. Examination on day 4 revealed significant cell death in wells containing HiDef and alfalfa-supplemented media. The suboptimal attachment of cells in serum-free media is often due to the absence of essential extracellular matrix proteins, such as fibronectin, laminin, and vitronectin, which are naturally present in serum and play a critical role in mediating cell adhesion through integrin binding. In serum-free conditions, the lack of these adhesion molecules necessitates the use of alternative surface coatings (*e.g*., recombinant proteins, and synthetic peptides) to mimic the extracellular matrix environment and facilitate proper integrin-mediated adhesion (Lam & Longaker, 2012; Li et al., 2020; Tragoonlugkana et al., 2024). Based on these findings, three different coatings—vitronectin, fibronectin, and laminin, each at a concentration of 1.5 µg/mL—were evaluated to determine their potential to support cell adhesion and proliferation in HiDef and alfalfa-supplemented media (Experiment II). Following coating, cells were counted, and doubling times were calculated on the third day of the subsequent phase. In both HiDef and alfalfa-supplemented media, the use of vitronectin reduced the doubling time by 3 hours compared to gelatin-coated controls, whereas no significant change was observed with Fibronectin or Laminin. Cells were then seeded for the following passage and observations on day four confirmed attachment to the wells. Microscopic examination indicated successful initial growth, predominantly with vitronectin coating. However, by day 6, before passage, cells began to exhibit morphological changes and detachment. The doubling time for cells coated with Vitronectin increased to 60 hours, while those coated with Fibronectin and Laminin reached 75 hours. Therefore, vitronectin demonstrated superior cell adhesion and growth compared to other coatings, aligning with findings from earlier studies by Stout et al. (2022, 2023). Stout et al. (2022) faced challenges with cell attachment in serum-free media containing recombinant albumin, and to overcome this issue, they tested various coatings, among them, vitronectin exhibited the best cell adhesion and growth (Stout et al., 2022).

Additionally, Stout et al. (2023) used vitronectin as the coating for developing serum-free media supplemented with rapeseed proteins. They found that this combination allowed for four passages (13 days), which is more than our developed media. The study also highlighted the importance of using optimal surface coatings to mimic the extracellular matrix, as the absence of crucial adhesion proteins like fibronectin, laminin, and vitronectin in serum-free environments often leads to suboptimal cell attachment (Stout et al., 2023).

These results underscore the critical role of vitronectin in enhancing cell adhesion and proliferation in specialized culture conditions. By providing a more supportive environment for integrin-mediated adhesion, vitronectin enables better cell growth and stability across multiple passages, making it a valuable component in the development of serum-free media for cultured meat production.

### 3.4 Metabolic profiling of fresh and spent media

Fresh and spent media analysis was conducted to track the metabolites and chemicals produced/consumed by the cells during the study. This approach will enable the optimization of culture media by revealing specific nutrient consumption patterns and the accumulation of metabolic byproducts. By understanding these dynamics, the media composition can be adjusted to better support cell growth, enhance productivity, and maintain optimal cell health throughout the culture process. Figure 3 shows the results of fresh and spent media analysis for OGM and protein isolates at different concentrations on day three.

**Figure 3.**
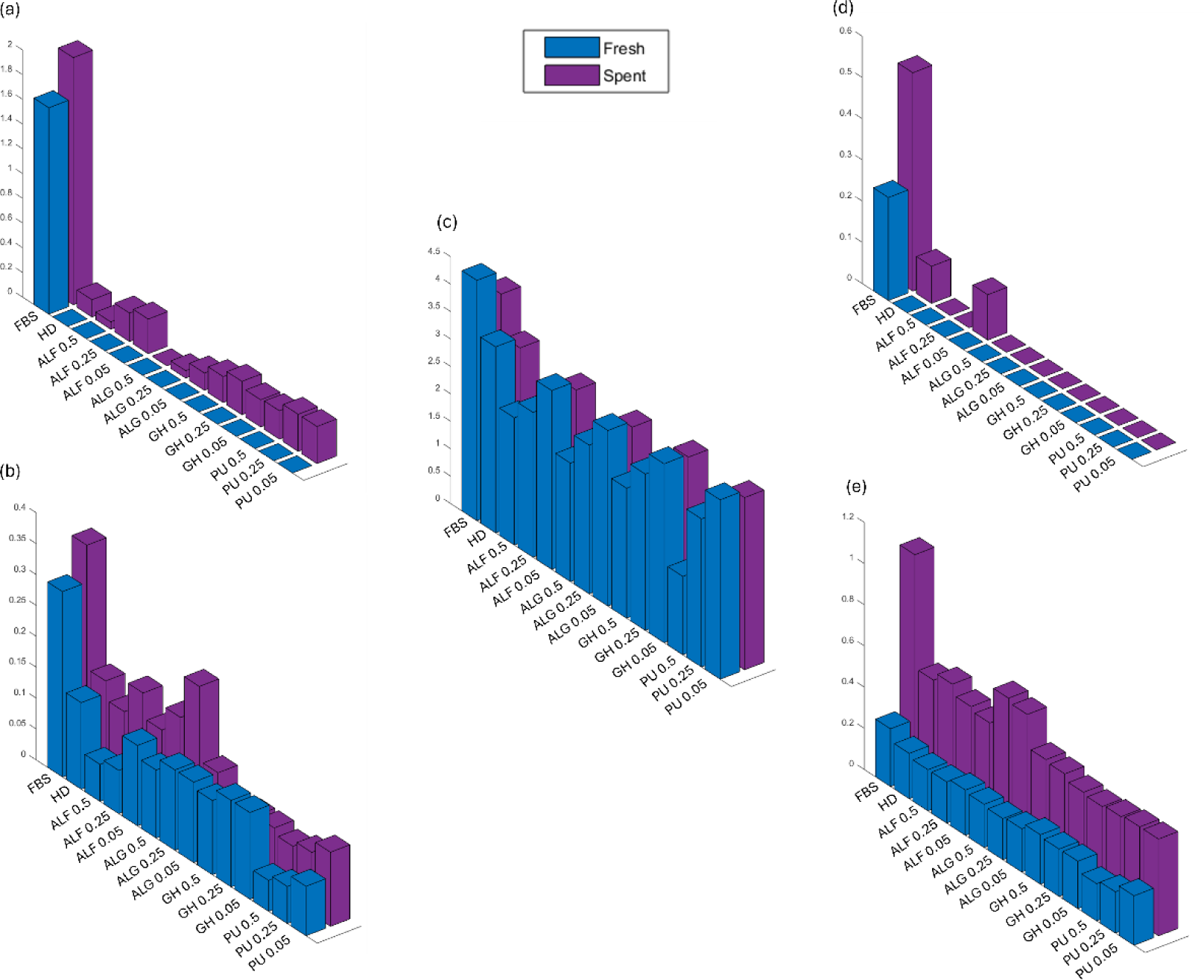
Analysis of fresh and spent media for different culture media and protein isolates at various concentrations on day three. The graphs represent the concentrations of key metabolites and chemicals, including (a) Glutamine, (mmol/L), (b) Glutamic acid (mmol/L), (c) Glucose (g/mL), (d) Lactate (g/mL), and (e) Ammonium (mmol/L) under both fresh and spent conditions. The x-axis categorizes the various media types, including FBS, HiDef, ALF, ALG, GH, and PU at different concentrations (0.05, 0.25, 0.5 mg/mL). The y-axis differentiates between fresh and spent media conditions. Only the alfalfa protein isolate showed lactate production in the spent media, indicating active cell metabolism, while other proteins did not.

Lactate production, an indicator of active cell metabolism, was observed exclusively in the spent media of alfalfa, whereas other protein isolates did not exhibit lactate production. Interestingly, alfalfa at lower concentrations showed a relatively low ammonium concentration compared to most of the other media. Ammonia is generated from the degradation of L-glutamine and other metabolic processes, and it can be toxic to cells by disrupting cellular metabolism, inhibiting ATP production, and acidifying the cytoplasm. The accumulation of these by-products poses challenges in large-scale cultured meat production, necessitating strategies for their removal or reduction to maintain a conducive environment for cell growth (Wang et al., 2018). Thus, despite the higher cell growth and increased metabolic activity observed, alfalfa exhibited lower levels of ammonia production by cells. This suggests that alfalfa not only supports cell proliferation but also minimizes the generation and accumulation of ammonia which could indicate more efficient nitrogen utilization, making alfalfa a promising supplement in cell culture media where controlling toxic byproducts is crucial for maintaining optimal cell conditions. This would also reduce the cost of operation significantly, making the cultivated meat economically viable.

Based on the analysis of fresh and spent media (Figure 3), glutamine in HiDef, was not detected. Glutamine is recognized as a primary energy source and a crucial precursor for nucleotide and protein synthesis (Yoo et al., 2020). This underscores the importance of media analysis for tracking various metabolites to identify and optimize key factors. For instance, O’Neill et al. (2022) highlighted the necessity of species- and cell-type-specific media optimization in cultivated meat production. Through spent media analysis of embryonic chicken muscle precursor cells, chicken fibroblasts, and murine C2C12 myoblasts, they demonstrated significant differences in nutrient utilization patterns, indicating that a universal medium formulation is unlikely to be effective across diverse cell types. Their findings validated the need for targeted media formulations to improve cost-effectiveness and efficiency in cultivated meat production, alongside novel approaches to streamline media optimization efforts (O’Neill et al., 2022).

To address the deficiency in glutamine and explore its impact on cell growth, a short-term experiment was conducted by supplementing the best concentration of alfalfa (0.05 mg/mL) with different concentrations of L-glutamine—specifically, 0.5, 1.6, and 3.0 mM. The objective was to assess the effects of different glutamine levels on cell growth and metabolism, with the goal of identifying the optimal concentration for enhancing cell proliferation and health. On the third day of the experiment, both fresh and spent culture media were analyzed, and proliferation assays were conducted using CyQuant (Figure 5). Figure 4 illustrates the results of the fresh and spent media following glutamine supplementation.

**Figure 4.**
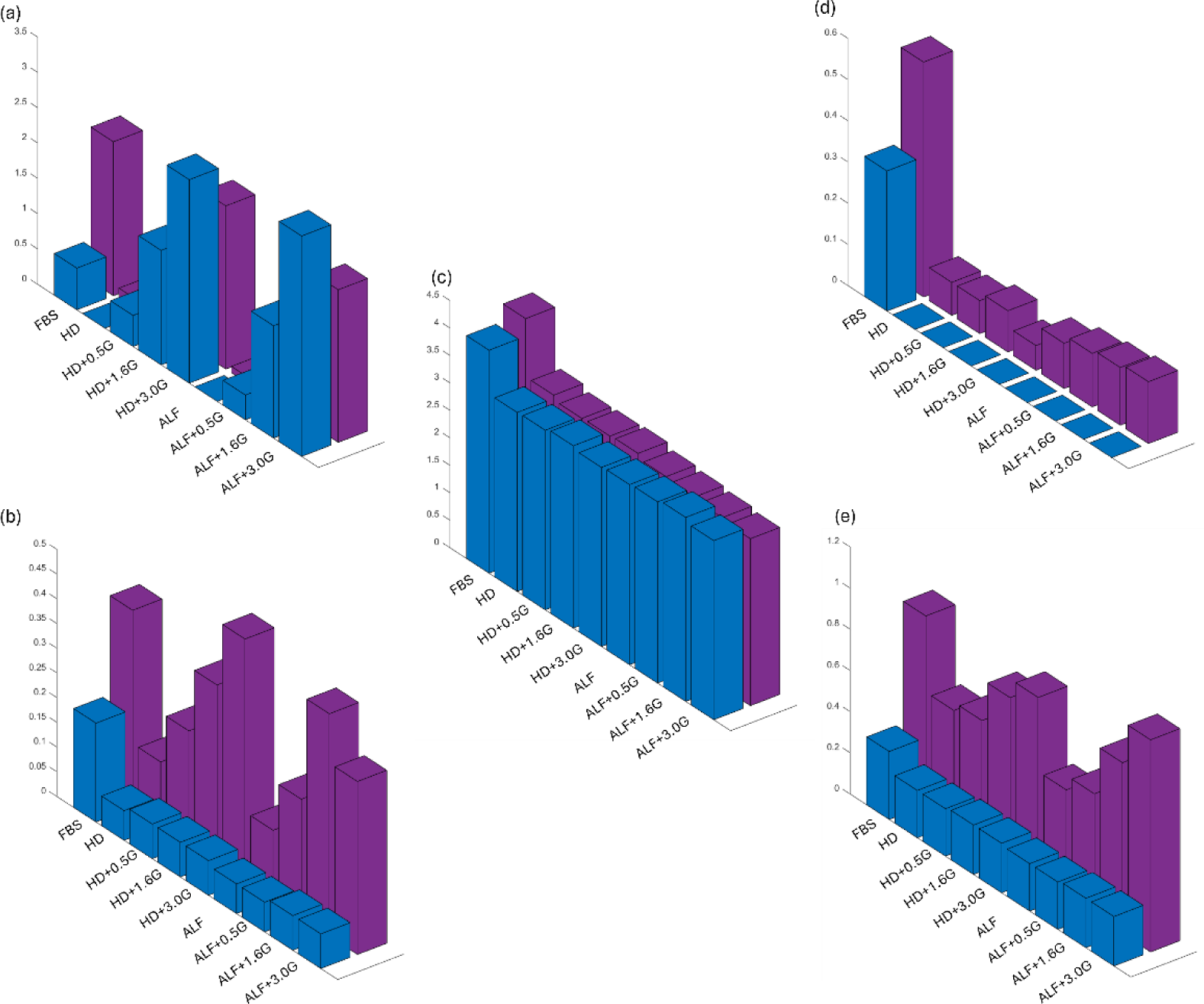
Analysis of fresh and spent media for different culture media on day three. HiDef (HD) and alfalfa (ALF) supplemented with various concentrations of L-glutamine. The graphs display the concentrations of key metabolites and chemicals, including (a) Glutamine (mmol/L), (b) Glutamic acid (mmol/L), (c) Glucose (g/mL), (d) Lactate (g/mL), and (e) Ammonium (mmol/L) under fresh and spent conditions. The x-axis categorizes the different media types, including FBS, HiDef, and alfalfa at a concentration of 0.05 mg/mL, with additional groups supplemented with 0.5, 1.6-, and 3.0-mM L-glutamine. The y-axis differentiates between fresh and spent media conditions. Notably, a correlation was observed between the concentration of added glutamine and ammonium production in the spent media, where increased glutamine levels led to a proportional rise in ammonium.

**Figure 5.**
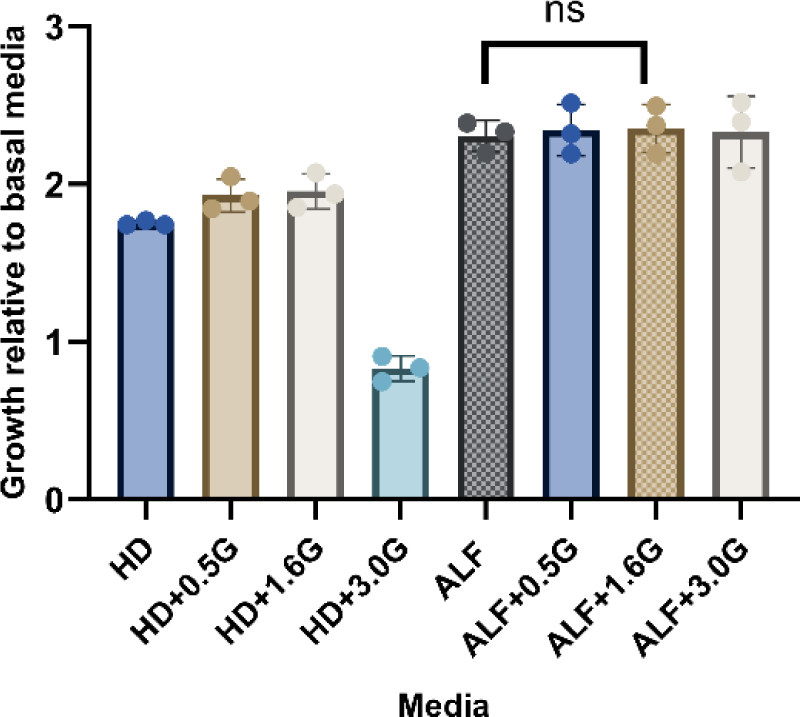
CyQUANT results from a three-day short-term growth study on the effect of glutamine supplementation. The experiment involved supplementing alfalfa and HiDef media with varying concentrations of L-glutamine (0.5, 1.6, and 3.0 mM) to evaluate the effect of glutamine on cell proliferation. Although glutamine supplementation appeared to enhance cell growth, the increase was not statistically significant compared to alfalfa without glutamine. Statistical significance was calculated by an unpaired t-test between alfalfa (A) 0.05mg/mL and alfalfa supplemented with 1.6 mM glutamine (G). Further long-term studies may provide more insights into the impact of glutamine on cell growth. (n=3 distinct samples)

A notable correlation was observed between the concentration of added glutamine and ammonium production in the spent media. As the concentration of glutamine increases, there is a proportional rise in ammonium levels. This relationship underscores the metabolic pathways involved in glutamine catabolism, particularly the deamination process where glutamine is converted to glutamate, releasing ammonium as a byproduct. This increase in ammonium production highlights a critical consideration in media formulation, as elevated ammonium levels can inhibit cell growth and function. Despite glutamine potential to enhance cell proliferation, the proliferation assay results (Figure 5) showed that this increase was not statistically significant when compared to the alfalfa media without glutamine supplementation. This suggests that while glutamine supplementation influences metabolic byproducts, its impact on cell growth may require further investigation, particularly over longer experimental durations to fully understand its effects on bovine satellite cell proliferation and health. In addition, these results indicate that alfalfa provided well-balanced amino acids for the cells, which extra concentrations of glutamine did not have any impact on cell proliferation.

## 4. Conclusion

This study highlights the potential of alfalfa protein isolates as a viable alternative to FBS in serum-free media for cultivating bovine satellite cells. Short-term growth studies indicated that low concentrations of alfalfa protein isolate significantly enhanced cell proliferation, closely matching the performance of FBS-supplemented media. While FBS contains numerous proteins essential for cell adhesion, none of the coatings used in this study supported long-term cell adhesion in serum-free media, indicating the need for further work to identify coatings or methods that support this requirement. Additionally, the glutamine supplementation experiment suggested a trend toward improved cell growth, though the results were not statistically significant, underscoring the need for further research to confirm its effectiveness and determine optimal conditions. These findings contribute to the development of sustainable and cost-effective cell culture media, essential for advancing the cultivated meat industry. Future research should focus on exploring additional protein isolates and optimizing basal media and essential cell growth factors to enhance cell growth and adhesion in serum-free conditions.

## Acknowledgments

This research was financially supported by the Agriculture and Food Research Initiative (AFRI) Sustainable Agricultural Systems program, grant no. 2021-699012-35978 from the USDA National Institute of Food and Agriculture, and Texas A&M AgriLife Research.

## Supplementary figures

**Figure S1:**
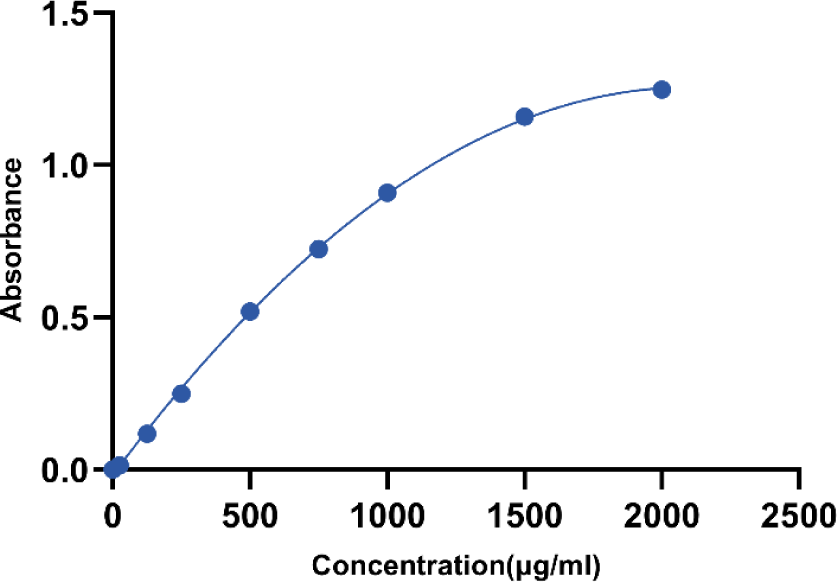
Bradford assay standard curve for protein quantification: The standard curve was generated using bovine serum albumin (BSA) standards ranging from 0 to 2000 µg/mL. Absorbance values at 595 nm were plotted against protein concentrations. Data points are the mean ± standard deviation of three independent replicates. A polynomial model was used to determine the protein isolate concentration.

**Figure S2:**
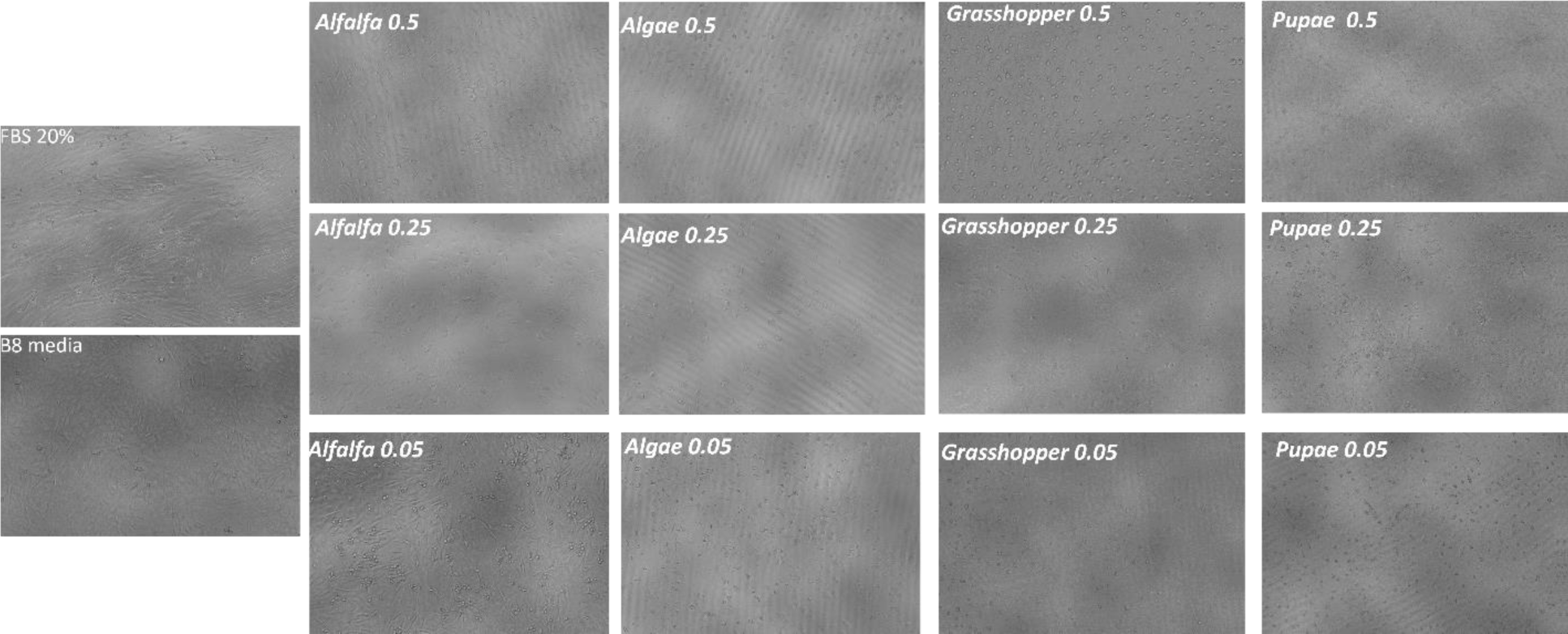
Brightfield images of OGM, HiDef and four protein isolates at various concentration on day three of growth; Algae, grasshopper, and silkworm pupae protein isolates at all concentrations, and alfalfa protein isolate at the highest concentration (0.5 mg/mL), showed poor cell growth and suboptimal cellular morphology. Alfalfa protein at the lowest concentration (0.05 mg/mL) demonstrated significant positive effects on cell growth and morphology, comparable to the control group with HiDef.

